# Latent Effector Capacity Governs Reversible T Cell Exhaustion: A Mathematical Model for Mechanistically Predictive AI in PD-1 Blockade

**DOI:** 10.64898/2026.04.13.717714

**Authors:** Aubrey Y. Liew, Ying Li, Haidong Dong

**Affiliations:** Department of Immunology, Mayo Clinic College of Medicine and Science, Rochester, MN; Department of Quantitative Health Sciences, Mayo Clinic, Jacksonville, FL; Department of Urology, Mayo Clinic, Rochester, MN

## Abstract

T cell exhaustion is commonly viewed as a terminal differentiation state marked by irreversible loss of effector function during chronic infection and cancer. However, the rapid restoration of cytotoxic activity following PD-1 checkpoint blockade challenges this view, revealing a central paradox: T cells that appear functionally inert can regain effector function on timescales incompatible with de novo differentiation or extensive epigenetic reprogramming. To resolve this contradiction, we present a mathematical framework that explicitly decouples latent effector capacity from active effector output. We define latent effector capacity as a slow, history-dependent state variable representing preserved epigenetic accessibility and regulatory readiness at effector loci, distinct from instantaneous transcriptional activity. Within this framework, PD-1 signaling functions as a reversible, graded masking mechanism that suppresses effector realization without erasing latent capacity, thereby explaining the coexistence of preserved chromatin accessibility, rapid functional rebound, and heterogeneous responses to checkpoint blockade. Incorporating nonlinear self-maintenance of epigenetic programs together with checkpoint-dependent erosion of latent capacity reveals a bistable regime and a history-dependent point of no return, beyond which exhaustion becomes irreversible. Critically, the model demonstrates that PD-1 checkpoint blockade unmasks pre-existing effector potential but cannot recreate lost capacity, because therapeutic reversibility is governed by the prior dynamical stability of a latent epigenetic state rather than by instantaneous transcriptional output. This framework establishes a mathematical foundation for mechanistically predictive AI in PD-1 blockade therapy by identifying latent, history-dependent variables that can be inferred from epigenetic and transcriptional data to predict therapeutic responsiveness and irreversibility.

## I. Introduction

A defining feature of chronic infection and cancer is the emergence of so-called *exhausted* T cells (Tex), a population characterized by diminished cytokine production, impaired cytotoxicity, sustained expression of inhibitory receptors such as PD-1, and altered transcriptional profiles ^1-3^. For many years, these cells were viewed as terminally dysfunctional, i.e. locked into an irreversible state of differentiation that precluded meaningful contribution to immunity against pathogens or cancer cells^4-6^. This view was grounded in observations of reduced effector gene expression, defective proliferative capacity, and stable transcriptional signatures associated with long-term antigen exposure^1,7,8^. However, the clinical success of immune checkpoint blockade has fundamentally challenged this interpretation. Remarkably, T cells that appear functionally inert immediately prior to therapy can rapidly regain effector activity following PD-1 blockade, exhibiting restored cytotoxic killing and clonal expansion on timescales shorter than those required for de novo differentiation or extensive epigenetic reprogramming in both preclinical and clinical studies^9-12^. The speed and robustness of this response are incompatible with models in which exhaustion reflects wholesale erasure of effector programs^9,13,14^. Instead, these observations imply that, at least in a substantial fraction of exhausted T cells, core components of effector identity remain preserved in a latent form^13,15-17^.

This paradox, *apparent terminal dysfunction followed by rapid functional recovery*, suggests that exhaustion cannot be fully explained by irreversible loss of differentiation state. Rather, it points to the existence of an active, reversible suppressive mechanism that constrains effector function without eliminating the underlying molecular infrastructure required to execute it^18,19^. Chromatin accessibility studies have shown that exhausted T cells often retain open regulatory regions at canonical effector loci^17,20^, including genes encoding cytotoxic molecules and inflammatory cytokines^21,22^. Similarly, lineage-defining transcription factors associated with effector identity can remain bound at key regulatory elements even when downstream gene expression is attenuated^8,23,24^. Together, these findings indicate that exhaustion involves dissociation between *competence* and *expression*^*25,26*^. To resolve this conceptual tension, we propose that T cell state is governed by two partially decoupled variables. The first is a latent effector capacity, reflecting the epigenetic, chromatin, and regulatory architecture accumulated through prior antigenic stimulation. This latent capacity encodes T cell readiness to mount an effector response via its accessible enhancers, poised promoters, and stabilized transcription factor networks, but without necessarily implying ongoing transcription or function. Importantly, this component evolves on relatively slow timescales, shaped by the history of T cell receptor engagement, inflammatory cues, and epigenetic reinforcement mechanisms, and is therefore resistant to rapid collapse.

The second variable is the active effector output, representing realized gene expression and functional execution at a given moment in time. This output is highly dynamic, sensitive to proximal signaling inputs, and subject to rapid suppression by inhibitory pathways. In this framework, immune checkpoint signaling, i.e. most prominently through PD-1, acts primarily on this active layer, attenuating transcriptional realization of effector programs downstream of preserved epigenetic competence. Crucially, PD-1 engagement masks effector activity without directly dismantling the chromatin landscape that supports it^27^. This decoupling provides a unifying explanation for several hallmark features of T cell exhaustion. It accounts for the coexistence of preserved epigenetic accessibility with diminished cytokine production^1,24^, explains why exhaustion can be rapidly reversed in some contexts yet refractory in others^26,28^, and clarifies why phenotype-based measurements alone can be misleading indicators of true functional potential of T cells^22^. Within this view, exhausted T cells are not uniformly inert^8,22,23^, but instead occupy a spectrum of states defined by how much latent effector capacity remains and how strongly that capacity is suppressed at the level of realization. By explicitly separating *capacity* from *output*, we shift the interpretation of exhaustion from a static terminal fate to a dynamic, history-dependent process. This perspective reframes immune checkpoint blockade not as a force that reprograms T cells de novo, but as an intervention that unmasks pre-existing potential, which is effective only to the extent that such potential has been preserved. Establishing this conceptual distinction is essential for understanding heterogeneity in therapeutic response, defining the point at which exhaustion becomes irreversible, and developing quantitative frameworks that integrate transcriptional, epigenetic, and functional data into a coherent model of T cell dysfunction.

Taken together, these considerations have important implications for predictive modeling of immune checkpoint therapy. Current efforts to predict response to PD-1 blockade largely rely on static molecular features or black-box machine-learning classifiers trained on outcome labels, approaches that implicitly assume instantaneous transcriptional states are sufficient to determine therapeutic responsiveness. However, the rapid yet heterogeneous responses observed following PD-1 blockade, together with the persistence of epigenetic “scars” in non-responding cells^6^, suggest that response is governed by latent, history-dependent cellular states rather than snapshot measurements alone. The conceptual separation between latent effector capacity and active effector output introduced here therefore naturally motivates a different class of predictive AI architectures—ones that explicitly represent latent biological state variables, reversible inhibitory masking, and cumulative signaling history, and evolve these quantities according to mechanistically informed dynamics. In this view, predictive AI is not an auxiliary analytic layer but a direct extension of the mathematical structure of exhaustion itself, providing a principled framework for integrating epigenetic and transcriptional data to forecast response, durability, and irreversibility of PD-1 blockade.

## II. Latent Effector Capacity as a Dynamical State Variable

To formalize the concept that exhausted T cells can retain hidden functional potential, we introduce latent effector capacity as an explicit dynamical variable. We define this quantity, denoted by the vector **E(t)**, as a measure of epigenetic accessibility and regulatory readiness at canonical effector loci, including *GZMB, PRF1*, and *IFNG*.

**E**(*t*) ∈ ℝ^*N*^: latent effector capacity vector (gene-specific chromatin and regulatory readiness)

Rather than reflecting instantaneous transcriptional activity, **E(t)** captures the *structural and regulatory state* of the chromatin landscape that permits rapid effector gene expression when inhibitory constraints are relieved. This distinction is critical, as numerous studies demonstrate preserved chromatin accessibility and transcription factor occupancy at effector regions in exhausted T cells^8,22^, indicating that effector identity can persist in a latent, masked form.

Biologically, latent effector capacity is accumulated during periods of productive T cell receptor (TCR) engagement through chromatin remodeling, enhancer activation, and stabilization of transcription factor networks downstream of strong antigenic stimulation^29,30^. These processes occur on relatively slow timescales and are known to imprint durable epigenetic features that outlast acute signaling events. Conversely, during chronic antigen exposure and sustained inhibitory signaling, this epigenetic infrastructure erodes gradually, reflecting progressive chromatin closing, loss of enhancer activity, and destabilization of effector-associated regulatory circuits^6,31^. Importantly, this decay is substantially slower than the suppression of active transcription, consistent with the observation that exhausted T cells can rapidly reacquire effector function following check-point blockade without requiring extensive reprogramming^9,10,14,32^.

We capture these biological processes using a minimal first-order dynamical equation for latent effector capacity:

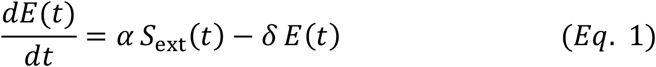

where ***S***_**ext**_(***t***) represents time-dependent external activation signals such as antigen availability, costimulatory cues, or inflammatory cytokines. The parameter ***α*** quantifies the efficiency with which these external inputs are converted into lasting epigenetic priming, integrating factors such as chromatin remodeler recruitment, histone modification kinetics, and transcription factor binding^29,33^. The decay constant ***δ*** represents slow exhaustion-associated silencing of effector loci driven by sustained inhibitory signaling, metabolic stress, and repressive chromatin modifications^34^.

This formulation embodies several key T cell biological principles^4,35,36^. First, epigenetic priming is integrative: transient activation signals incrementally load latent capacity over time, rather than triggering binary state transitions. Second, epigenetic decay is slow and proportional to existing capacity, reflecting the progressive nature of exhaustion rather than abrupt loss of effector identity. Finally, the linear structure of Eq. 1 emphasizes that latent effector capacity evolves independently of instantaneous gene expression, allowing it to remain elevated even when transcriptional output is strongly suppressed by checkpoint signaling.

As a consequence, Eq. 1 naturally explains how exhausted T cells can occupy a state characterized by high latent potential but low functional output. When *S*_ext_(*t*) has been historically strong, *E*(*t*) may remain elevated for prolonged periods, even if active effector gene expression is masked by inhibitory pathways such as PD-1 signaling. This decoupling establishes the foundation for rapid functional rebound upon checkpoint blockade: effector activity can re-emerge on transcriptional timescales because the underlying epigenetic substrate has already been established. Thus, modeling latent effector capacity as a dynamical state variable provides a mechanistic bridge between stimulation history, epigenetic memory, and reversible T cell dysfunction.

## III. PD-1 Signaling as a Masking (Rheostat) Function

A Hill-type masking function describes how increasing amounts of a “masker” gradually and non-linearly reduce the influence of another signal with a tunable sensitivity and steepness. Based on our previous conceptual mode of T cell responses^37^, we model PD-1–mediated checkpoint inhibition using a Hill-type masking function to reflect the graded, saturable, and reversible nature of PD-1 signaling on effector gene expression. Biologically, PD-1 engagement does not erase effector programs but instead suppresses their transcriptional realization through cooperative, multi-step inhibition of TCR and costimulatory signaling^38-41^. A Hill formulation captures this behavior by introducing a tunable inhibitory threshold and sensitivity, allowing effector output to transition smoothly from near-uninhibited to strongly suppressed as PD-1 signaling intensity increases. Importantly, this masking term acts multiplicatively on effector expression without directly altering latent effector capacity, formalizing the conceptual distinction between preserved epigenetic potential and actively functional output. In this framework, PD-1 blockade corresponds to removal of the masking constraint, enabling rapid functional rebound from pre-existing latent capacity without requiring de novo reprogramming.

We represent checkpoint inhibition using a Hill-type masking function:

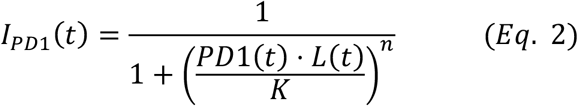

where ***I***_***PD*1**_(***t***) denotes the effective PD-1 inhibitory input experienced by a T cell at time *t*. It represents the *instantaneous strength of checkpoint signaling* that acts upstream of transcriptional and functional suppression of effector programs. ***PD*1**(***t***) is receptor expression, ***L***(***t***) is ligand availability (e.g., PD-L1 or PD-L2). The parameter ***K*** defines the effective inhibitory threshold of PD-1 signaling, integrating downstream signaling efficiency and transcriptional susceptibility, and sets how much PD-1–PD-L1 engagement is required to mask otherwise preserved latent effector capacity. Importantly, *K* should not be interpreted as the biochemical dissociation constant of PD-1–PD-L1 binding. Instead, it is a *composite phenomenological parameter* that aggregates multiple downstream processes between receptor engagement and transcriptional suppression. As PD-1 blockade effectively reduces the numerator PD1 ⋅ *L* in cells with high *K* cross below the inhibitory threshold rapidly leads to strong rebound, while in cells with low *K* may remain masked despite blockade unless ligand is nearly eliminated. To that end, progenitor-like exhausted T cells (Tpex) may have higher *K*, while terminally exhausted T cells (Ttex) would have lower *K*. This provides a mechanistic explanation for why cells with comparable PD-1 expression can respond very differently to checkpoint blockade^42,43^.

The Hill coefficient ***n*** parameterizes the effective cooperativity and sharpness of PD-1–mediated suppression of effector gene expression. Low values of *n* (∼ 1) correspond to gradual, proportional inhibition as PD-1 signaling increases, consistent with weakly cooperative or diffuse check-point control. Higher values of *n* produce a steeper, threshold-like transition in effector output, reflecting the convergence of multiple inhibitory steps downstream of PD-1 engagement and capturing switch-like masking behavior characteristic of deeper exhaustion states. Importantly, *n* does not imply direct molecular cooperativity at the receptor level; rather, it serves as a phenomenological measure of how decisively checkpoint signaling suppresses transcriptional realization of otherwise intact effector programs.

Together, as illustrated in Figure 1, *K* shifts the masking curve left or right along the PD-1·PD-L1 axis; while *n* determines whether masking is gradual or threshold-like. Thus, they define whether PD-1 acts as a dimmer or a binary gate on effector realization. Therefore, a Hill-type masking function models PD-1 signaling as a tunable, reversible dimmer switch that suppresses effector expression in proportion to inhibitory signal strength, without erasing the underlying effector potential.

**Figure 1.**
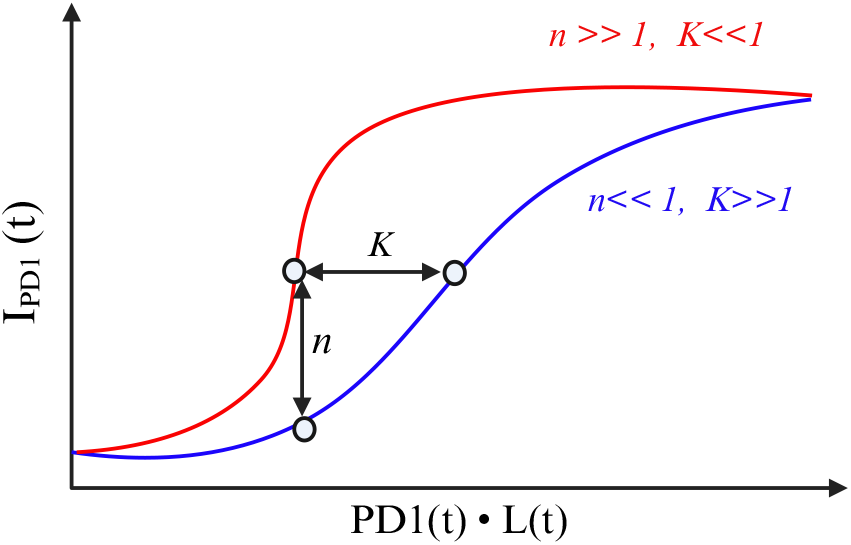
PD-1 masking function and parameter interpretation. Hill-type masking function *I*_PD1_(*t*) (Eq. 2) illustrating how checkpoint signaling suppresses effector realization as a function of effective PD-1 engagement, PD1(*t*) ⋅ *L*(*t*). The parameter *K* sets the inhibitory threshold, determining sensitivity to PD-1 signaling, while the Hill coefficient *n* controls the steepness of masking, ranging from graded rheostat-like suppression (*n*≪1) to sharp, threshold-like gating (*n*≫1). Together, *K* and *n* define whether PD-1 acts as a tunable dimmer or a switch-like gate on effector output without altering latent effector capacity. The graph is created with BioRender.

## IV. Active Effector Gene Expression Dynamics

We denote the vector of actively expressed effector genes by **G**(*t*):

**G**(*t*) ∈ ℝ^*N*^: vector of actively expressed effector genes

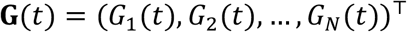

With genes explicitly resolved, the dynamics of actively expressed effector genes become:

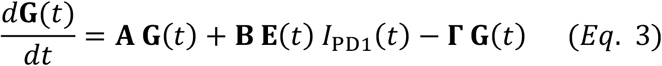

In this equation, **A** encodes cross-regulation, co-expression, and feed-forward reinforcement among effector genes (e.g., IFNG–TBX21 coupling, cytotoxic module coherence).

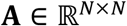

We use **B** for the recruitment / realization matrix (typically diagonal) to map latent capacity into transcriptional output when inhibition is relieved:

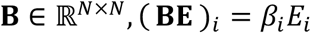

The PD-1 masking term (Equation 2) acts multiplicatively but does not alter **E**(*t*), while it suppresses realization of functional outputs. Additionally, we introduce a decay matrix **Γ** that accounts for gene-specific mRNA/protein turnover.

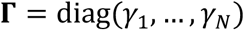

For each individual effector gene *i*, we will have:

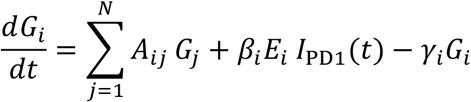

This highlights that the effector capacity (*E*_*i*_) is permissive, but not deterministic. The checkpoint PD-1 signaling acts upstream of transcriptional realization, not epigenetic identity. At the mean-time, different effector genes can decay, reinforce, or rebound at different rates.

Equation 3 describes the dynamical regulation of active effector gene expression as a balance between transcriptional production, checkpoint-mediated suppression, and molecular decay. The production term summarizes intrinsic gene–gene regulatory interactions and activation-dependent transcriptional inputs that would drive effector expression in the absence of inhibitory signaling. PD-1 signaling enters as a multiplicative masking term in the denominator, progressively attenuating effective transcriptional output as checkpoint engagement increases. This formulation reflects the biological observation that PD-1 suppresses the realization of effector outputs rather than eliminating the underlying regulatory or epigenetic machinery^44^.

The decay term accounts for turnover of effector transcripts and proteins, ensuring that expression levels represent a dynamic steady state rather than a permanently fixed fate. Therefore, low effector gene expression in exhausted T cells emerges from an actively maintained equilibrium between synthesis and checkpoint suppression, not from irreversible loss of effector identity^45^. Upon PD-1 blockade, removal of the masking constraint allows effector expression to rapidly increase, driven by pre-existing latent capacity rather than de novo reprogramming.

## V. Functional History: The Integral Representation

Instantaneous measures of effector gene expression capture only the current transcriptional state of a T cell, but immune function is inherently cumulative. Cytokine secretion, cytotoxic killing, and tissue remodeling integrate effector activity over time and cannot be inferred from a single snapshot. To formally distinguish *latent or instantaneous effector potential* from *realized immune function output*, we therefore introduce an explicit integral representation of effector output. This formulation allows us to encode the *functional history* of the cell and resolve the paradox that exhausted T cells can retain high latent potential while exhibiting little accumulated functional impact.

We define the realized effector state *G*(*T*) as the time integral of instantaneous effector gene expression dynamics:

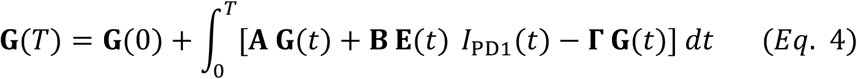

Here, **G**(*T*) ∈ ℝ^*N*^ represents a cumulative realized effector output vector, integrating functional execution over time ***t***, with **G**(0): denoting the initial effector baseline. **E**(*t*) ∈ ℝ^*N*^: latent effector capacity (epigenetic and regulatory readiness). Each term in the integrand has a direct biological interpretation:

### Gene–gene regulatory reinforcement (A ⋅ G(*t*))

This term captures the internal regulatory architecture of effector gene programs, representing gene–gene interactions that reinforce, coordinate, and stabilize effector identity once transcription has been initiated. The matrix **A** encodes cross-regulatory influences among effector loci, including positive feedback loops, co-expression modules, and mutual reinforcement mediated by shared transcription factors and signaling pathways. Biologically, these interactions reflect well-characterized effector circuits such as the coupling between IFNG and TBX21, the coordinated activation of cytotoxic genes including GZMB, PRF1, and NKG7, and broader transcriptional modules that promote coherent effector function. In the absence of checkpoint inhibition, these intrinsic dynamics can sustain or amplify effector gene expression over time, enabling self-reinforcing transcriptional states that persist even as upstream activation signals fluctuate. Within the model, this term formalizes the idea that effector programs possess internal stability and structure, allowing active effector expression to be maintained through network-level reinforcement rather than relying solely on continuous external stimulation. Consequently, gene–gene regulatory reinforcement provides a mechanistic basis for persistence, coherence, and robustness of effector identity once latent capacity has been successfully realized.

### Latent-to-active realization under checkpoint masking B E(*t*) *I*_PD1_(*t*)

This term represents the conversion of preserved epigenetic and regulatory potential into observable effector gene expression, conditional on the strength of inhibitory checkpoint PD-1 signaling. The latent effector capacity **E**(*t*) encodes gene-specific chromatin accessibility, poised enhancers, and stabilized transcription factor networks that enable rapid transcriptional activation. The recruitment matrix **B** maps this latent capacity into transcriptional output, specifying the efficiency with which each effector locus can be realized once inhibitory constraints are relieved. Critically, immune checkpoint signaling through PD-1 regulates this conversion process via the multiplicative masking function *I*_PD1_(*t*), which attenuates transcriptional realization without directly altering the underlying epigenetic state. As a result, effector output can be strongly suppressed even when substantial latent capacity is retained. This formulation formalizes exhaustion as a state of *masked competence*: checkpoint signaling constrains gene expression at the level of realization rather than erasing effector identity, thereby allowing rapid functional rebound when the masking constraint is removed while preserving the possibility of irreversible failure only when latent capacity itself decays.

### Decay term −**Γ*G***(***t***)

Accounts for molecular turnover of effector transcripts and proteins, ensuring that effector expression reflects an actively maintained balance rather than a permanently fixed fate.

### Biological implication of the integral formulation in Eq. 4

It fundamentally reframes PD-1 biology by explicitly separating instantaneous transcriptional suppression from cumulative immune function. In this framework, PD-1 does not merely reduce effector gene expression at a given moment, but persistently attenuates the *integrand* of functional output over time. As a result, exhaustion emerges not from a single decisive failure of effector differentiation, but from prolonged integration of effector programs under sustained inhibitory masking. Even when latent effector capacity **E**(*t*) remains high, chronic PD-1 signaling keeps the realization term **BE**(*t*)*I*_PD1_(*t*) suppressed at each time point, leading to minimal accumulation of **G**(*T*). Thus, exhausted T cells can appear functionally inert despite retaining substantial epigenetic and regulatory readiness, resolving the paradox between preserved molecular competence and poor cumulative immune performance.

This formulation highlights *functional history* as an essential dimension of T cell state. Instantaneous measurements of effector gene expression capture only the current value of **G**(*t*), whereas immune efficacy—such as cytotoxic killing, cytokine delivery, and tissue remodeling—depends on the *area under the curve* accumulated over time (functional history). Chronic PD-1 signaling flattens this curve by maintaining prolonged suppression, thereby limiting functional impact even when short-term rebound potential exists^39,46^. Exhaustion is therefore recast as a history-dependent process: low cumulative function reflects prolonged exposure to inhibitory signaling rather than irrevocable loss of effector identity.

Within this context, PD-1 blockade acquires a precise mechanistic interpretation. Removing PD-1– mediated masking increases the integrand immediately, allowing rapid growth of **G**(*T*) driven by pre-existing latent capacity rather than slow epigenetic reprogramming. The speed of functional rebound following PD-1 blockade is thus explained naturally by the release of an already-loaded effector program^14,16,47^, rather than by de novo differentiation. Conversely, the model clarifies why PD-1 blockade fails in terminally exhausted cells: if latent capacity **E**(*t*) has already decayed due to cumulative inhibitory exposure, increasing the integrand has little effect because the substrate for integration has collapsed.

## VI. Mapping Latent Capacity to Realized Function

A central outcome of our framework is a formal, mechanistic link between preserved epigenetic potential and observed effector behavior. While earlier sections separate latent effector capacity from actively expressed effector genes, immune function ultimately depends on how these two components interact. We therefore propose an explicit mapping from latent capacity to realized effector expression that resolves the apparent contradiction between preserved molecular potential and functional silencing in exhausted T cells.

We define instantaneous active effector expression as

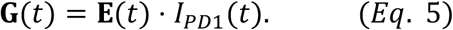

where *E*(*t*) denotes the latent effector capacity and *I*_PD1_(*t*) represents a PD-1–dependent inhibitory masking function bounded between 0 and 1.

This formulation embodies two key biological assumptions. First, latent effector capacity is permissive rather than deterministic: accessible chromatin, poised transcription factors, and preserved regulatory architecture enable effector gene expression but do not guarantee it. Second, checkpoint signaling regulates realization rather than identity: PD-1 engagement does not erase effector programs but instead suppresses their transcriptional deployment through reversible signaling mechanisms. Expressing effector output as a product of capacity and masking captures this gating relationship in the minimal mathematical form. Importantly, the multiplicative structure implies that either term can independently constrain effector output. If latent capacity is lost (*E*(*t*) → 0), effector expression necessarily collapses, corresponding to terminal exhaustion. Conversely, if inhibitory signaling is strong (*I*_PD1_(*t*) → 0), effector expression remains low even when substantial latent capacity is preserved.

Under chronic antigen stimulation, our model predicts a characteristic regime in which *E*(*t*) remains elevated due to accumulated epigenetic priming, while sustained PD-1 signaling drives *I*_PD1_(*t*) to low values. In this regime,

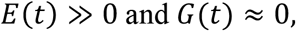

yielding the exhausted phenotype: *high potential, low output*. This relationship provides a mechanistic explanation for several hallmark observations of exhausted T cells. Molecular profiling often reveals preserved chromatin accessibility and transcription factor binding at effector loci, despite minimal cytokine production or cytotoxic activity. In the present framework, such cells occupy a state where epigenetic capacity is maintained but functionally masked by inhibitory signaling, rather than irreversibly erased.

Equation 5 also makes explicit why PD-1 blockade can induce rapid functional recovery. Because *E*(*t*) evolves on slow timescales governed by chromatin remodeling, its value at the time of intervention largely reflects prior stimulation history. Removal of checkpoint signaling increases *I*_PD1_(*t*) toward unity without requiring changes in *E*(*t*), immediately unmasking pre-existing effector capacity and allowing *G*(*t*) to rise sharply. Thus, rebound effector activity emerges naturally as a *state-dependent unmasking phenomenon*, rather than as de novo differentiation. This prediction aligns with the rapid kinetics of functional restoration observed experimentally following PD-1 blockade. By explicitly separating *capacity* from *realization*, Eq. 5 cautions against interpreting low effector gene expression as evidence of lost differentiation status. Instead, a low *G*(*t*) value may reflect either diminished epigenetic competence or active suppressive signaling, with profoundly different therapeutic implications. This distinction becomes critical when integrating transcriptional, epigenetic, and functional assays, as it explains why exhausted T cells can appear inert by expression-based metrics while retaining substantial latent responsiveness.

In summary, mapping latent effector capacity to realized function via a multiplicative checkpoint-masking relationship provides a unifying explanation for reversible dysfunction, functional rebound, and the hidden potential of exhausted T cells. This linkage establishes the foundation for subsequent analysis of irreversibility, cumulative functional output, and the point at which epigenetic capacity itself collapses.

## VII. PD-1 Blockade and the Rebound Effect

A hallmark feature of exhausted T cells (or at least a subset of them) is their capacity for rapid functional restoration following immune checkpoint blockade, despite minimal effector activity immediately prior to treatment^10,11,25^. Within the latent effector capacity framework developed above, this phenomenon emerges naturally as a direct consequence of removing inhibitory masking from a pre-existing reservoir of epigenetically encoded potential.

We model PD-1 blockade by setting ligand availability to zero,

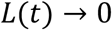

corresponding biologically to therapeutic interruption of PD-1–PD-L1 interactions. In the full model, PD-1 signaling suppresses the realization of effector programs through a Hill-type masking term that appears in the denominator of the effector expression function. Eliminating ligand availability collapses this inhibitory term, thereby removing checkpoint-mediated suppression without altering the underlying latent effector capacity.

Formally, in the limit of complete checkpoint blockade, instantaneous effector expression obeys

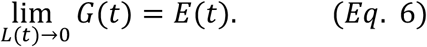

This identity states that, once inhibitory signaling is removed, active effector gene expression directly reflects the latent effector capacity already encoded in the cell. Importantly, Eq. 6 makes no assumptions about changes in chromatin state, transcription factor availability, or regulatory architecture at the time of treatment. Instead, effector output increases because the epigenetic and regulatory groundwork was established earlier during antigen exposure (functional history) and subsequently preserved in a masked form.

This relationship provides a precise mathematical definition of functional rebound or resilience^13,15,18,19^. Prior to blockade, exhausted T cells may exhibit negligible effector gene expression due to strong PD-1–dependent suppression, even while maintaining substantial latent capacity. Upon blockade, the masking constraint is instantaneously lifted, allowing effector expression to rise on timescales governed by transcription and translation rather than slow epigenetic remodeling. As a result, functional restoration occurs rapidly, consistent with experimental observations showing swift re-emergence of cytokine production and cytotoxic activity following PD-1 inhibition^13,19^.

Crucially, Eq. 6 clarifies that rebound does not require reversal of exhaustion-associated chromatin changes or de novo differentiation into effector states at the time of therapy. Instead, rebound reflects state-dependent unmasking: the conversion of preserved latent capacity into observable function once inhibitory signaling is removed. This distinction explains how exhausted T cells can appear irreversibly dysfunctional by transcriptional or functional measures immediately before treatment, yet respond robustly within hours to days following checkpoint blockade. Furthermore, the magnitude of rebound predicted by Eq. 6 is determined entirely by the value of the latent effector capacity *E*(*t*) at the time of intervention. Cells with high preserved capacity exhibit strong functional recovery, whereas cells in which *E*(*t*) has already decayed—corresponding to terminal exhaustion—show minimal or no rebound even when PD-1 signaling is fully blocked. Thus, PD-1 blockade unmasks, but cannot recreate, effector potential. In this way, Eq. 6 provides a concise mechanistic explanation for therapeutic heterogeneity observed in immune checkpoint therapy. Functional rebound is not an intrinsic property of PD-1 blockade alone, but rather a consequence of the interaction between blockade and the pre-existing epigenetic state of responding T cells. The equation therefore links molecular state, treatment timing, and therapeutic outcome within a single, experimentally interpretable framework.

## VIII. The Point of No Return: Loss of Epigenetic Plasticity

While earlier sections describe exhaustion as a reversible state arising from inhibitory masking of preserved latent effector capacity, prolonged chronic stimulation ultimately leads to a qualitatively different regime in which the latent epigenetic capacity itself deteriorates^6,23,27^. Experimentally, this terminally exhausted state is characterized by stable loss of chromatin accessibility at effector loci (epigenetic scar)^6^, reduced transcription factor binding, and failure to respond to checkpoint blockade^27^. To capture this transition within our framework, we extend the latent capacity dynamics to explicitly model self-maintenance of epigenetic programs and their checkpoint-dependent destabilization.

We modify Eq. 1 by introducing a nonlinear self-maintenance term and a checkpoint-dependent decay term, yielding

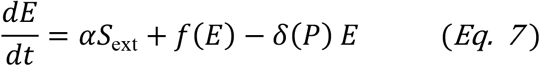

Here, *E* denotes the latent effector capacity. The external stimulation term *αS*_ext_ captures antigen-driven loading of epigenetic potential, as before. The newly introduced function ***f***(***E***) represents autonomous, self-reinforcing maintenance of accessibility and regulatory architecture, while ***δ***(***P***)***E*** represents active erosion of capacity driven by persistent check-point signaling. Biologically, these added terms formalize the observation that chromatin states are not purely passive memories of past stimulation. Once established, accessibility at effector loci can be maintained by positive feedback involving lineage-defining transcription factors, enhancer– promoter looping, and cooperative chromatin remodeling. Conversely, sustained inhibitory signaling through PD-1 and related pathways promotes repressive chromatin modifications and destabilization of effector programs. Equation 7 therefore models latent capacity as the outcome of a competition between self-maintenance and checkpoint-driven decay.

From there, we further model self-maintenance using a saturating positive feedback function,

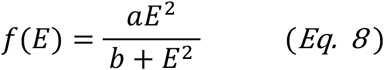

This Hill-type form captures two essential properties of epigenetic reinforcement. First, feedback is weak at low capacity, reflecting the inability of sparse or fragmented regulatory elements to stabilize themselves. Second, once capacity exceeds a critical level, feedback becomes strong and saturating, enabling robust maintenance of effector-associated chromatin states. Importantly, this function introduces nonlinearity into the dynamics of *E*, allowing for multiple steady states.

As a result, the system can exhibit *bistability*: a high-*E* state corresponding to preserved epigenetic plasticity, and a low-*E* state corresponding to terminal exhaustion. The existence of these states depends on the balance between feedback strength ***a***, external stimulation, and the effective decay rate imposed by PD-1 signaling. ***b*** sets the value of *E*^2^ at which the positive-feedback function reaches half of its maximum, and determines the horizontal position of the feedback curve along the *E*-axis. i.e. Small *b* → feedback turns on at low *E*, i.e. effector programs are easy to stabilize → favors reversibility and memory-like behavior; Large *b* → feedback requires high *E* to become effective, i.e. stabilization requires strong, sustained priming → favors collapse into terminal exhaustion. Thus, parameter *b* sets the threshold for self-reinforcing epigenetic maintenance, defining the amount of latent effector capacity required before positive feedback stabilizes chromatin accessibility and preventing collapse into terminal exhaustion.

The transition between reversible and irreversible exhaustion occurs when the high-*E* steady state is lost. Mathematically, this corresponds to a saddle-node bifurcation, defined by the condition:

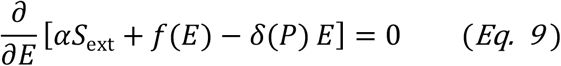

At this critical point, the stable high-capacity equilibrium collides with an unstable equilibrium and disappears. Beyond this threshold, no amount of external stimulation or removal of checkpoint inhibition can restore a high-*E* state, because the self-maintenance machinery required to stabilize epigenetic accessibility has already collapsed. Biologically, this bifurcation defines a *Point of No Return*: a history-dependent threshold after which effector chromatin states are no longer recoverable. Prior to this point, PD-1 blockade primarily reduces effective decay by lowering *δ*(*P*), allowing the system to relax back to the high-capacity attractor. Beyond it, the attractor itself no longer exists.

A central implication of this analysis is that PD-1 blockade unmasks but does not recreate epigenetic capacity. In the reversible regime, lowering *δ*(*P*) shifts the system back into the basin of attraction of the high-*E* state, leading to rapid functional rebound. In contrast, once the saddle-node bifurcation has occurred, removal of checkpoint signaling has little effect because the structural determinants of effector identity have been irreversibly lost. This framework explains why terminally exhausted T cells fail to respond to checkpoint therapy despite effective receptor blockade and intact proximal signaling. Failure of rescue does not reflect insufficient drug activity, but rather the absence of a latent epigenetic scaffold that can support effector gene expression. The model therefore provides a mechanistic basis for therapeutic timing effects, emphasizing that checkpoint blockade is most effective when applied *before* epigenetic plasticity crosses the point of no return.

## IX. Integral Criterion for Irreversibility

While the preceding sections define exhaustion as a dynamical competition between preserved latent effector capacity and checkpoint-mediated masking, an important unresolved question is *when* reversibility is lost altogether. Empirically, exhausted T cells exhibit a continuum of responses to PD-1 blockade, ranging from rapid functional rebound to complete refractoriness. These observations suggest that irreversibility is not determined by instantaneous signaling strength alone, but instead reflects the history of inhibitory exposure experienced by the cell.

To formalize this concept, we express the loss of epigenetic plasticity as an accumulated exposure criterion, in which irreversible exhaustion emerges once the time-integrated burden of PD-1 signaling exceeds a critical threshold:

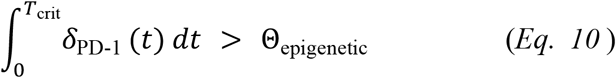

Here, ***δ***_**PD-1**_(***t***) represents the instantaneous effective strength of PD-1–mediated inhibitory signaling, incorporating receptor expression, ligand availability, and downstream signal propagation. The upper limit ***T***_**crit**_ denotes the time at which the system crosses from a reversible into an irreversible regime. The threshold **Θ**_**epigenetic**_ represents a chromatin locking threshold: a quantitative measure of cumulative inhibitory exposure beyond which effector-associated epigenetic architectures can no longer be stably maintained. This integral formulation captures a central biological principle: epigenetic deterioration is driven by sustained signaling, not transient events. Brief or intermittent PD-1 engagement may suppress effector gene expression acutely, yet leave the underlying chromatin landscape intact. In contrast, prolonged and persistent checkpoint signaling gradually promotes repressive chromatin modifications, erosion of enhancer accessibility, and loss of reinforcing transcription factor occupancy. These processes are slow, cumulative, and inherently history dependent, making them poorly described by instantaneous threshold models.

Equation 10 therefore distinguishes two qualitatively different exhaustion regimes. In the reversible regime, the accumulated PD-1 signal remains below Θ_epigenetic_. In this case, latent effector capacity may be actively suppressed but is structurally preserved, allowing PD-1 blockade to rapidly unmask effector programs. In the irreversible regime, the accumulated inhibitory exposure surpasses the chromatin locking threshold, corresponding to stable loss of effector-associated epigenetic features. Beyond this point, removal of checkpoint signaling cannot restore function, as the latent epigenetic scaffold required for effector gene expression has collapsed. Importantly, Eq. 10 provides an equivalent, history-based description of the saddle-node bifurcation identified in the latent capacity dynamics. Whereas the bifurcation analysis characterizes irreversibility in terms of the disappearance of a high-capacity steady state, the integral criterion reframes this transition as a time-integrated damage process. The two descriptions are mathematically consistent: sustained PD-1 signaling both increases the cumulative integral in Eq. 10 and drives the system toward the loss of the high-capacity attractor. This perspective highlights that the timing of intervention is as critical as its strength. PD-1 blockade is effective when applied before the accumulated inhibitory exposure exceeds Θ_epigenetic_, at which point the system can relax back toward a high-capacity state. Once this threshold has been crossed, blockade reduces ongoing inhibitory signaling but cannot undo the historical accumulation that has already destabilized chromatin architecture.

Biologically, Θ_epigenetic_ can be interpreted as the cumulative level of repression required to lock effector loci into a stably inaccessible state. This may correspond to acquisition of repressive histone marks, DNA methylation, loss of enhancer–promoter looping, or depletion of key lineage-defining transcription factors. Different T cell populations, differentiation states, or tissue environments may exhibit distinct values of Θ_epigenetic_, providing a mechanistic explanation for heterogeneity in responsiveness to checkpoint therapy. From an experimental standpoint, Eq. 10 predicts that markers reflecting integrated inhibitory history, such as cumulative PD-1 engagement, duration of antigen exposure, or progressive chromatin closure measured by longitudinal ATAC-seq— will be stronger predictors of therapeutic reversibility than instantaneous PD-1 expression or signaling alone. This framework also suggests that early intervention, or strategies that intermittently relieve inhibitory signaling, may prevent cells from crossing the chromatin locking threshold altogether.

Taken together, the above considerations indicate that T-cell exhaustion can be succinctly expressed as a multiplicative relationship between a slow, history-dependent latent state and a rapidly reversible inhibitory constraint. In particular, instantaneous effector activity can be written as:

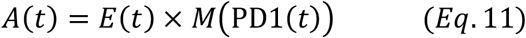

where ***E***(***t***) denotes latent effector capacity, reflecting the preserved epigenetic and regulatory architecture accumulated over prior stimulation history, and ***M***(**PD1**(***t***)) represents checkpoint-dependent masking that suppresses the realization of this capacity at a given moment in time. Within this formulation, PD-1 blockade acts primarily on the masking term and therefore restores effector activity only to the extent that latent capacity remains intact. Loss of reversibility arises when sustained inhibitory exposure destabilizes epigenetic self-maintenance, eliminating the high-*E* attractor and rendering increases in *M*(PD1) insufficient to recover function. The schematic graph illustrates this relationship (Figure 2), emphasizing how reversible suppression of effector output and irreversible loss of latent capacity together define the point of no return in T-cell exhaustion.

**Figure 2.**
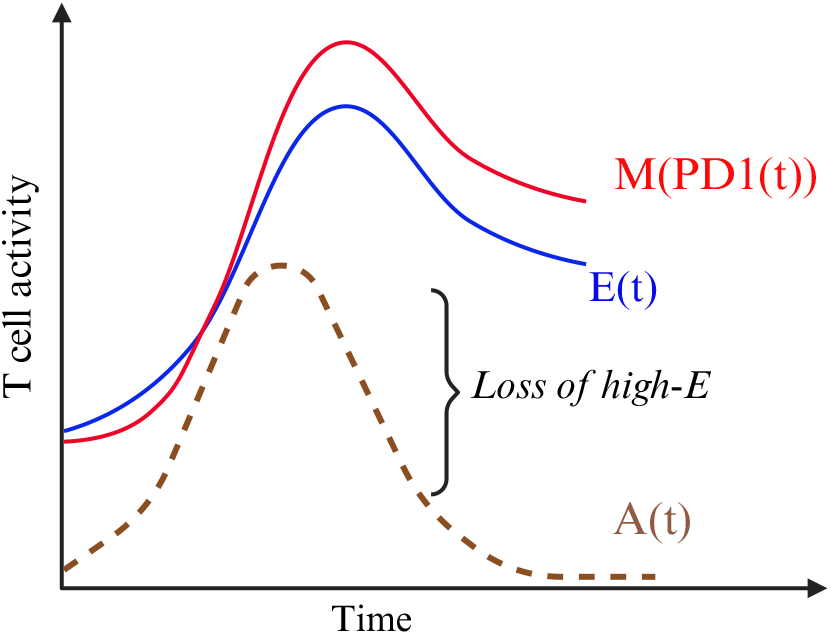
Conceptual schematic of the core equation governing T-cell exhaustion. Latent effector capacity *E*(*t*) represents the slow, history-dependent epigenetic and regulatory competence of effector programs, while checkpoint-dependent masking *M*(PD1(*t*)) suppresses the realization of this capacity without directly erasing it. Observed effector activity *A*(*t*) (dashed) arises as the product of E(t) x M(PD1(t)), explaining how T cells can retain substantial latent potential despite low functional output. Prolonged inhibitory signaling progressively destabilizes the high-*E* state; once epigenetic self-maintenance is lost, the high-capacity attractor disappears, defining a point of no return beyond which PD-1 blockade can no longer restore effector function. Time is shown schematically; irreversibility reflects a structural transition in latent state space rather than an acute signaling event. The graph is created with BioRender.

## X. Future direction: A Mechanistically Constrained AI Framework for Predicting Response to PD-1 Blockade

A central implication of the latent effector capacity framework developed here is that therapeutic response to PD-1 blockade is governed not by instantaneous transcriptional state alone, but by a set of *hidden, history-dependent dynamical variables*, including preserved epigenetic capacity, reversible inhibitory masking, and cumulative checkpoint exposure. These quantities are not directly observable, yet they are in principle inferable from multimodal molecular data. This perspective naturally motivates the development of mechanistically constrained artificial intelligence (AI) models that explicitly represent and evolve these latent state variables, rather than attempting to predict clinical response using black-box classifiers. In short, a correct AI to predict PD-1 therapy response is a mechanistically constrained Neural State-Space / Neural ODE model that learns latent effector capacity (E), checkpoint masking (I_PD1), and cumulative inhibitory history directly from epigenetic and transcriptional data, rather than predicting response labels directly.

### Mechanistic requirements for an AI predictor of checkpoint response

The mathematical model presented in this work imposes several solid structural requirements on any predictive framework for PD-1 blockade response. (1) the model explicitly separates latent effector capacity—a slow, epigenetically encoded variable—from active effector output, which is rapidly modulated by inhibitory signaling. (2) PD-1 is shown to act primarily as a *reversible masking mechanism* on effector realization, rather than as a driver of immediate epigenetic erasure. (3) irreversibility emerges only after time-integrated inhibitory exposure exceeds a critical threshold, corresponding to a loss of epigenetic self-maintenance. Together, these principles imply that patient response cannot be inferred from static measurements alone, but must be modeled as the outcome of a dynamical system with memory and bifurcation structure. Consequently, the appropriate AI architecture is not a conventional supervised classifier trained on response labels, but rather a *latent state–space model* that (i) infers unobserved biological state variables from molecular data, (ii) evolves these variables according to mechanistically informed dynamics, and (iii) uses the resulting trajectories to anticipate therapeutic outcomes under PD-1 blockade.

### Inference of latent effector capacity from epigenetic data

Within this framework, latent effector capacity serves as a foundational state variable encoding the preserved regulatory and epigenetic infrastructure of effector programs. Future AI models can infer this quantity directly from chromatin accessibility data, such as single-cell or bulk ATAC-seq, by learning structured representations of enhancer–promoter accessibility, transcription factor motif usage, and regulatory network coherence at effector gene loci. Importantly, this inferred latent state is not intended to represent current transcriptional activity, but rather the *potential* for rapid effector gene realization when inhibitory constraints are relieved, consistent with the distinction formalized in the present model. Encoding epigenetic data into a latent capacity variable also enables direct testing of key model predictions, including the existence of bistability, self-reinforcing epigenetic maintenance, and collapse beyond a point of no return. In this sense, AI-based inference of latent capacity provides a principled bridge between high-dimensional chromatin data and the low-dimensional dynamical variables required for predictive modeling.

### Modeling checkpoint masking and transcriptional realization

In parallel, transcriptional data can be used to infer the active effector state, representing realized gene expression at a given moment in time. Within the present framework, this state evolves rapidly and is multiplicatively gated by inhibitory checkpoint signaling. AI models that explicitly encode this multiplicative relationship can distinguish between low effector output arising from strong inhibitory masking versus true loss of epigenetic competence: two scenarios with fundamentally different therapeutic implications. Critically, modeling PD-1 signaling as a learned, graded masking function enables patient-specific inference of inhibitory sensitivity, rather than assuming uniform checkpoint effects across individuals^48^. Such an approach is naturally aligned with the Hill-type masking formulation introduced here and provides a mechanistic explanation for heterogeneous responses among patients with comparable PD-1 expression.

### Learning immune history and irreversibility

Perhaps the most distinctive contribution of the present framework is the formalization of *immune history* as a determinant of irreversibility. The integral criterion defining the transition to terminal exhaustion implies that the *cumulative burden* of inhibitory signaling, rather than its instantaneous magnitude, governs loss of responsiveness to PD-1 blockade. AI models that explicitly maintain and update a history-dependent internal state, analogous to an accumulated damage or exposure variable, are therefore uniquely suited to predict not only whether a patient will respond, but *when intervention has already become too late*. By embedding this cumulative exposure variable within a dynamical system that exhibits bifurcation behavior, AI predictors can output clinically interpretable quantities such as rebound potential, durability of response, and probability of irreversibility. These outputs move beyond binary response prediction and instead align with the mechanistic logic of the model.

### Toward predictive and interpretable digital immune models

Taken together, these considerations point toward a new class of *mechanistically grounded AI models* for immunotherapy: models that infer latent biological state variables from epigenetic and transcriptional data, evolve those states according to experimentally motivated dynamical equations, and simulate the consequences of therapeutic perturbations such as PD-1 blockade. In contrast to purely data-driven approaches, this architecture offers intrinsic interpretability, causal consistency, and the ability to perform counterfactual simulations, such as predicting the effect of earlier intervention or combination therapies aimed at preserving epigenetic capacity. In the long term, such models may serve as computational representations of *virtual T cells* or *digital immune twins*, integrating molecular state, signaling history, and epigenetic memory to forecast immune trajectories under chronic disease and therapy. Within this vision, latent effector capacity emerges not only as a conceptual advance, but as a practical organizing variable for predictive, quantitative immunology.

## XI. Limitations and Scope

While the present work establishes a mechanistically grounded mathematical foundation for reversible and irreversible T cell exhaustion, several limitations should be acknowledged. Importantly, these limitations primarily reflect deliberate modeling choices that prioritize interpretability, analytical tractability, and conceptual clarity, rather than empirical shortcomings. Where relevant, each limitation is supported by existing experimental evidence and directly motivates the AI-based extensions outlined in the above section.

### Model abstraction and molecular resolution

First, the framework intentionally adopts a coarse-grained representation of complex molecular processes, minimizing chromatin remodeling, transcription factor dynamics, metabolic constraints, and signaling cascades into a small number of effective state variables. Latent effector capacity is treated as a continuous scalar or low-dimensional vector, whereas epigenetic accessibility and regulatory readiness are known to be distributed across thousands of loci and organized into partially independent regulatory modules^5,6,23,29^. This abstraction sacrifices locus-specific granularity in favor of a unifying dynamical description capable of explaining rapid reversibility, preserved chromatin accessibility, and heterogeneous therapeutic responses across systems. Importantly, this simplification should not be interpreted as implying that effector programs are monolithic. Genome-wide profiling has demonstrated that cytotoxicity, cytokine production, proliferative capacity, and memory persistence can be epigenetically uncoupled during exhaustion^8,22,36^. Resolving these components will require integration of multi-modal single-cell epigenomic datasets, which naturally motivates higher-dimensional extensions of the present framework through mechanistically constrained AI models.

### Reduced representation of checkpoint networks

Second, the model focuses primarily on PD-1–mediated inhibition as the dominant reversible masking mechanism governing effector realization. While PD-1 signaling has been shown to suppress transcriptional and cytoskeletal programs without immediate erasure of effector chromatin states^27,38,44^, additional inhibitory receptors, including TIM-3, LAG-3, TIGIT, and suppressive cytokine pathways, also contribute to functional dysfunction in exhausted T cells^1,26,31,48^. These pathways are not explicitly modeled here and are instead subsumed into phenomenological parameters governing masking strength and decay rates. This reductionist choice enhances mathematical clarity but limits direct attribution of exhaustion dynamics to individual checkpoint pathways. Nonetheless, experimental studies support the conceptual validity of modeling inhibitory signaling as a composite masking effect rather than discrete binary switches^35,43^. The AI-based latent state-space framework proposed in future directions offers a principled mechanism for learning these composite inhibitory influences directly from data while preserving mechanistic interpretability.

### Cell-intrinsic focus and population-level effects

Third, the framework is explicitly cell-intrinsic and does not incorporate population-level processes such as clonal selection, cell-cell interaction/regulation, step-wise differentiation, or competitive expansion. In vivo responses to PD-1 blockade frequently involve selective expansion of progenitor-like exhausted T cell subsets with preserved epigenetic capacity, rather than direct reinvigoration of terminally exhausted cells^9,23,32^. By focusing on single-cell dynamical states, the present model does not attempt to disentangle reinvigoration from clonal replacement^9,49,50^. This limitation is intentional: the goal is to define the *intrinsic cellular conditions required for reversibility*, independent of downstream population dynamics. Coupling the present framework to population-level (spatial biology) or evolutionary models (cell development) represents an important future direction and will be necessary for linking single-cell trajectories to bulk clinical outcomes^9,14,16^.

### Parameter identifiability and empirical calibration

Fourth, while the mathematical formulation is mechanistically motivated and consistent with extensive transcriptional and epigenetic datasets, several parameters, particularly those governing epigenetic self-maintenance, check-point-dependent decay, and the chromatin locking threshold—are not directly measurable with current experimental tools. Experimental studies demonstrate durable epigenetic “scars” following chronic antigen exposure^6,27^, but quantitative thresholds governing the loss of reversibility remain poorly defined. Thus, our model should be interpreted as a *structural and predictive framework* rather than a fully parameterized simulator. This limitation underscores the need for longitudinal, perturbation-resolved datasets combining chromatin accessibility, transcriptional output, signaling history, and functional measurements. It also motivates the integration of mechanistically constrained AI approaches capable of inferring latent state variables and cumulative inhibitory exposure directly from high-dimensional data rather than relying on manually specified parameters.

### Scope of generalization

Finally, although this framework is developed in the context of CD8^+^ T cell exhaustion under chronic antigen stimulation and PD-1 blockade, similar separations between latent competence and realized function have been reported in T cell anergy, tolerance, memory differentiation, and immune resilience. Extending the model to these contexts will require careful biological validation and may necessitate additional state variables or alternative dynamical regimes.

### Perspective

Taken together, these limitations define the intended scope of the present model rather than diminishing its contribution. The framework establishes latent effector capacity as a central organizing variable and formalizes exhaustion as a history-dependent dynamical process with a mathematically defined point of no return, consistent with current transcriptional and epigenetic evidence^5,6,27^. The AI-based extensions introduced in the above section are not ancillary additions, but a natural continuation of this logic, enabling scalable inference, empirical grounding, and predictive application without abandoning mechanistic interpretability.

## Acknowledgments

We are grateful to members of the Dong laboratory and to our research and clinical collaborators for valuable discussions and for their contributions to the review of the relevant literature. This work was supported in part by Schmidt Sciences through funding for AI Research and Innovation and by the Mayo Clinic Comprehensive Cancer Center. Due to space limitations, we may not have cited all outstanding studies in the rapidly evolving field of T-cell exhaustion, and we acknowledge these contributions with appreciation. The authors declare no competing interests.

## Notes

### Competing Interest Statement

The authors have declared no competing interest.

